# In Vivo Two-Photon Imaging of Neuronal and Vascular Responses in Mice Chronically Exposed to Ethanol

**DOI:** 10.1101/471763

**Authors:** Phillip O’Herron, Phillip M. Summers, Andy Y. Shih, Prakash Kara, John J. Woodward

## Abstract

The effects of ethanol on brain function have been extensively studied using a variety of in vitro and in vivo techniques. For example, electrophysiological studies using brain slices from rodents and nonhuman primates have demonstrated that acute and chronic exposure to ethanol alters the intrinsic excitability and synaptic signaling of neurons within cortical and sub-cortical areas of the brain. In humans, neuroimaging studies reveal alterations in measures of brain activation and connectivity in subjects with alcohol use disorders. While complementary, these methods are inherently limited due to issues related to either disruption of normal sensory input (in vitro slice studies) or resolution (whole brain imaging). In the present study, we used 2-photon laser scanning microscopy in intact animals to assess the impact of chronic ethanol exposure on sensory evoked neuronal and vascular responses. Adult male C57BL/6J mice were exposed to 4 weekly cycles of chronic intermittent ethanol (CIE) exposure while control mice were exposed to air. After withdrawal (≤ 72 hr), a cranial window was placed over the primary visual cortex (V1) and sensory evoked responses were monitored using the calcium indicator OGB-1. CIE exposure produced small but significant changes in response amplitude (decrease) and orientation selectivity of V1 neurons (increase). While arteriole diameter did not differ between control and CIE mice under baseline conditions, sensory-evoked dilation was enhanced in vessels from CIE exposed mice as compared to controls. This was accompanied by a reduced latency in response to stimulation. In separate experiments, pial arteriole diameter was measured in the barrel cortex of control and CIE exposed mice. Baseline diameter of barrel cortex arterioles was similar between control and CIE exposed mice but unlike vessels in V1, sensory-evoked dilation of barrel cortex arterioles was similar between the two groups. Together the results of these studies suggest that chronic exposure to alcohol induces changes in neurovascular coupling that are region dependent.

## Introduction

Alcohol abuse and alcoholism are major contributors to mortality and morbidity and impose a significant sociological and economic burden on the US population (2013). Understanding the neurobiological consequences that result from excessive drinking and that contribute to relapse is thus an important goal of alcohol research. Current thinking suggests that alcohol-dependent individuals have difficulty in gaining voluntary control over their drinking and this is associated with deficits in cortical-based cognitive ability and alterations in visual and other sensory processing when presented with alcohol-related cues. Growing evidence also suggests that neurons within the visual cortex are involved in reward processing similar to those in other areas such as the ventral tegmental area, amygdala and frontal cortex. For example, neurons in the primary visual cortex of rats that receive a reward paired with a visual stimulus fire in a way that predicts when the reward will be delivered (Shuler and Bear, 2006). These findings are complemented by studies in humans that show that visual cortical activity in addicts but not controls, is enhanced by visual cues related to drug use (Yalachkov *et al*, 2010). This enhanced responsiveness to drug-related cues (attentional bias) can persist for long periods of time even in abstinent individuals and may contribute to relapse by triggering strong feelings of craving for the drug.

Chronic alcoholism is also associated with deficits in cerebrovascular function and impairments in activity-dependent recruitment of blood flow (termed *neurovascular coupling*) may contribute to the disruption of normal brain function that persists during protracted periods of relapse (Hanlon *et al*, 2014). Understanding the cellular and molecular processes that underlie these deficits may lead to novel treatment strategies that help restore control over excessive drinking and prevent relapse.

A major limitation in the study of these processes is the lack of suitable experimental models and approaches that allow for in vivo analysis of brain function at the cellular and network level. While results from whole-brain imaging and EEG recording studies have identified significant impairments in brain function in alcoholics, these methods suffer from a lack of precision especially with respect to cellular subtype. At the other extreme, electrophysiological analysis of neuronal activity using in vitro slice preparations allows for detailed analysis of the electrical and signaling properties of single cells but has limited translational capability and low throughput. In this study, we use two-photon microscopy and a well-validated mouse model of ethanol dependence to investigate the effects of chronic ethanol exposure on neuronal and vascular function in vivo.

## Methods

### Animals and Chronic Ethanol Exposure

Male C57/BL6J mice were obtained from Jackson Laboratories (Bar Harbor, ME) (https://www.jax.org/strain/013636) at 9 weeks of age. They were group-housed (4/cage) and allowed to acclimatize to the colony room for at least one week in a temperature and humidity controlled AAALAC-approved facility. Animals were maintained on a 12-hour light/dark cycle with lights off at 09:00 am and had ad libitum access to food and water. All animals were treated in strict accordance with the NIH Guide for the Care and Use of Laboratory Animals and all experimental methods were approved by the Medical University of South Carolina’s Institutional Animal Care and Use committee.

### Chronic intermittent ethanol exposure

After acclimatization, mice were treated with repeated cycles of chronic intermittent ethanol (CIE) vapor exposure, a protocol that induces dependence in mice (Lopez and Becker, 2005). Briefly, each day for 4 days, mouse cages containing bedding, food and water bottles were placed in vapor inhalation chambers and exposed for 16 hrs to either air (control) or air saturated with ethanol vapor (CIE group). Air flow in both chambers was maintained at 5 L/min. At the end of each daily exposure, fresh bedding, food and water was added and cages were returned to the housing colony for 8 hr. At the end of the fourth day, mice underwent 3 days of abstinence before beginning the next weekly cycle of ethanol exposure with a total of four weeks of ethanol or air exposure. Ethanol (95%) was volatized by passing air through a submerged air stone and the resulting vapor was mixed with fresh air and delivered to Plexiglas inhalation chambers to maintain consistent ethanol concentrations between 17-21 mg/L air in the chamber. This yielded blood ethanol concentrations (BEC) in the range of 225-300 mg/dl. Prior to entry into the ethanol chambers, CIE mice were injected intraperitoneally (20 ml/kg body weight) with ethanol (1.6 g/kg; 8% w/v) and the alcohol dehydrogenase inhibitor pyrazole (1 mmol/kg) to maintain stable blood ethanol levels (Nimitvilai *et al*, 2016). Air control mice were similarly handled but were injected with saline and pyrazole before being placed in air inhalation chambers. Chamber ethanol concentrations were monitored daily and air flow was adjusted to maintain concentrations within the specified range. In addition, blood samples were collected and determined from all animals to monitor BECs during the course of inhalation exposure. Average BECs during the 4 weeks of CIE exposure was 290.49 ± 27.32 mg/dl. Mice were used for in vivo imaging studies 3-7 days following the final vapor exposure.

### Two-photon imaging

#### Visual Cortex

The protocol for the visual cortex imaging and analysis studies are described in detail in previous reports (O’Herron *et al*, 2016; O’Herron *et al*, 2012; O’Herron *et al*, 2018). Briefly, mice were anaesthetized with a mixture of fentanyl citrate (0.04–0.05 mg/kg), midazolam (4–5 mg/kg), and dexmedetomidine (0.20–0.25 mg/kg) and craniotomies (2–3 mm square) were opened over the primary visual cortex. The calcium indicator Oregon Green 488 Bapta-1 AM (OGB-1 AM) and a red dye (Alexa 633 or Alexa 594) for visualization were mixed in a glass pipette and injected into the craniotomy with air pressure puffs. After a 1 hr dye loading period, the dura was removed and the craniotomies were sealed with agarose (1.5–2% dissolved in artificial cerebrospinal fluid) and covered with a 5-mm glass coverslip. The concentrations of the anesthetics were lowered (fentanyl citrate: 0.02–0.03 mg/kg/hr, midazolam: 1.50–2.50 mg/kg/h, and dexmedetomidine: 0.10–0.25 mg/kg/hr) during two-photon imaging. Fluorescence was monitored with a custom-built microscope (Prairie Technologies) coupled with a Mai Tai (Newport Spectra-Physics) mode-locked Ti:sapphire laser (810 nm or 920 nm) with DeepSee dispersion compensation. Excitation light was focused by a 40X (NA 0.8, Olympus) water immersion objective. Full frame imaging of approximately 300 μm square windows was obtained at approximately 0.8 Hz. Drifting square-wave grating stimuli were presented on a 17-inch LCD monitor and were presented at 16 directions of motion in 22.5° steps for 6.5 seconds with 13 seconds of blank before each stimulus. Each condition was repeated at least 8 times except for two runs with only 5 repetitions.

#### Sensorimotor Cortex

Studies of whisker-evoked vascular responses were performed on awake mice habituated to head-fixation, as previously described (Summers *et al,* 2017). Briefly, mice underwent surgery for placement of a cranial window over the left sensorimotor cortex and a mounting plate over the right cortex. After recovery, mice were habituated to head fixation for 2-7 days prior to imaging. During each imaging session, mice were briefly anesthetized with 4% MAC isoflurane followed by an infraorbital vein injection of 2 MDa fluorescein-dextran (FD2000S; Sigma-Aldrich; prepared at a concentration of 5% (w/v) in sterile saline). After waking, they were allowed to recover for 30 min prior to imaging. A Sutter Moveable Objective Microscope and a Coherent Ultra II Ti:Sapphire laser source (800 nm excitation) were used to collect movies encompassing 312 by 244 μm areas of the pial surface at a frame rate of 4 Hz. Arteriole diameter was measured offline using full-width-at-half-maximum calculations of the fluorescence intensity profile across the vessel width. To evoke sensory-dependent responses, whiskers in head-fixed, awake mice were stimulated with air puffs (8 Hz, 20 ms pulse, 10 s pulse train, 35 p.s.i. from air tank) directed at the mystacial pad contralateral to the hemisphere with the imaging window. The air stream was placed 2 cm from the whiskers and focused on rows B to D. A second air puffer was directed at the tail as a control for general arousal. Whisker and tail stimulation trials were presented in random order during the experiment. Ten trials of each stimulation type were collected for each imaged location. Each trial consisted of a 30 s baseline, 10 s stimulation, and 50 s post-stimulation period.

### Statistical Analysis

Results were analyzed for statistical significance using Prism 7.0 software (GraphPad, La Jolla, CA). Data are presented as mean ± SEM and were considered significantly different when p < 0.05.

## Results

In the first study, neural and vascular responses were monitored in the primary visual cortex (V1) of mice exposed to 4 cycles of CIE or air. Examples of calcium signals from individual mouse V1 neurons during presentation of stimuli are shown in Figure 1A. For neural responses, we calculated response amplitudes expressed as the percent change from baseline for each neuron’s preferred stimulus direction. As shown in Figure 1B, stimulus-evoked responses from air treated mice averaged ~13% and this response was slightly reduced (mean difference −0.89 ± 0.002) in CIE exposed mice. Due to the large number of neurons sampled, this difference, although small, was statistically significant (unpaired t-test, t=506.3, df=5420, p<0.0001).

**Figure 1.**
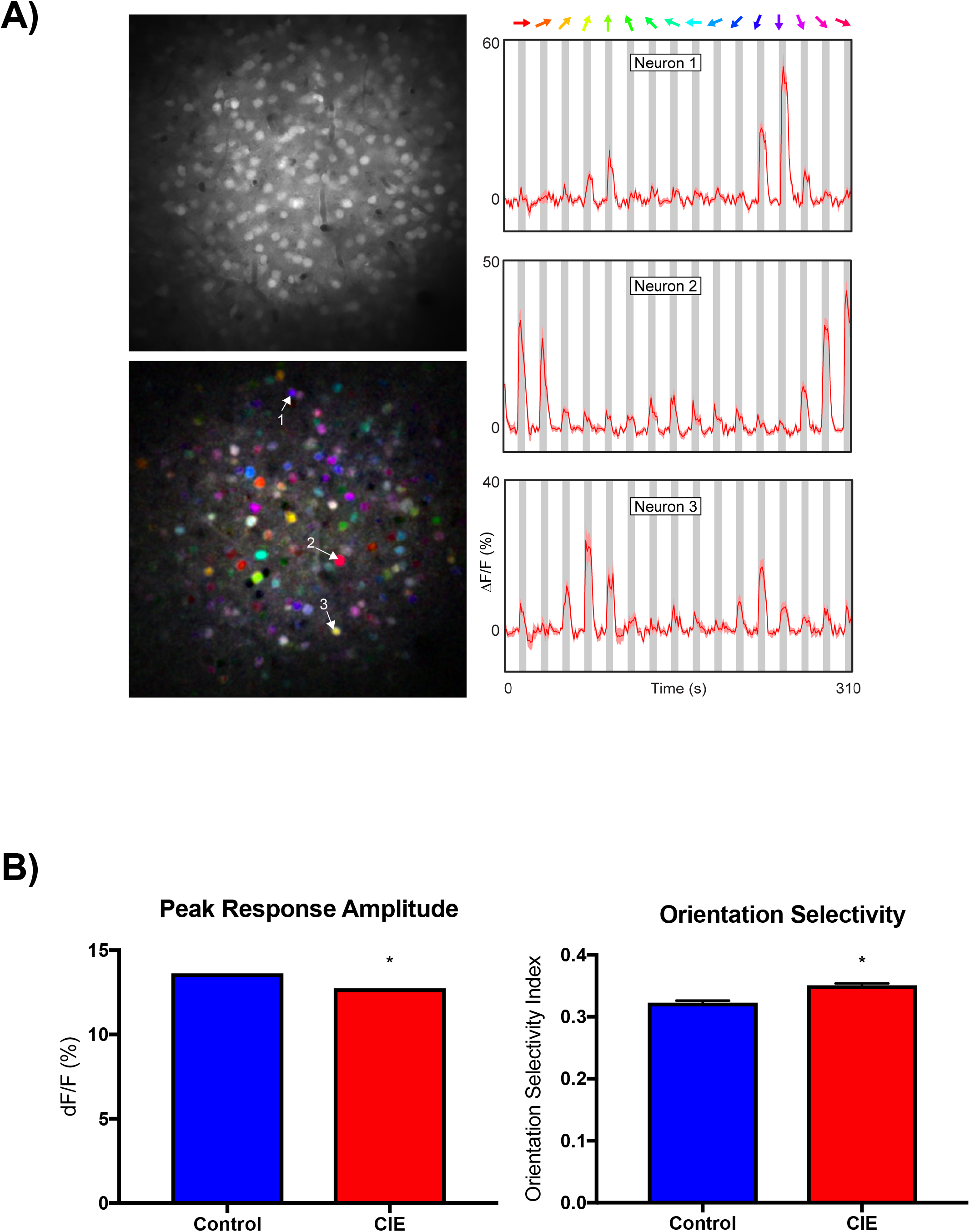
Effects of CIE exposure on sensory-evoked responses in visual cortex neurons. A) Right column: representative examples of neuronal calcium signals during presentation of drifting grating stimuli (colored arrows). Left column: gray scale image of field of view (top) and pixel-based image map of stimulus direction (bottom). The hue corresponds to the preferred direction, the saturation indicates selectivity and the brightness gives the response strength. Numbered arrows indicate neurons shown in corresponding time series graphs. B) Graphs show mean (± SEM) response amplitude (left) and orientation selectivity index (right) for control and CIE treated mice. Symbol (*): p < 0.05; unpaired t-test.

In addition to amplitude responses, we also measured the tuning characteristics of visually responsive neurons in V1 cortex expressed as the orientation selectivity index (OSI). In Air mice, OSI averaged approximately 0.3 (Figure 1B) and we observed a slight (mean difference 0.028 ± 0.004) but statistically significant increase (unpaired t-test, t=6.42, df=5420, p<0.0001) in this value in CIE treated mice. In the same animals, we used Alexa 633 to measure blood vessel diameter under baseline conditions and during presentation of the visual stimuli (see Figure 2A for examples of these responses). Baseline V1 vessel diameter was not different between air and CIE treated mice (Figure 2B). During visual stimulation, V1 vessels from both air and CIE treated mice dilated with those in CIE mice showing a significantly greater increase (Figure 2C; unpaired t-test, t=2.204, df=40, p<0.03). To assess whether CIE treatment altered the response characteristics of vessels, we calculated the response latency of vessel dilation expressed as the time for the response to reach 2 or 3 times the standard deviation of the baseline diameter. As shown in Figure 2D, latencies in CIE treated mice were significantly shorter than those in air controls (unpaired t-test 3SD, t=2.354, df=40, p<0.02; 2SD, t=2.507, df=40, p<0.02). Fitting a linear regression to the rising phase of vessel dilation revealed the same effect (unpaired t-test, t=2.392, df=40, p<0.02).

**Figure 2.**
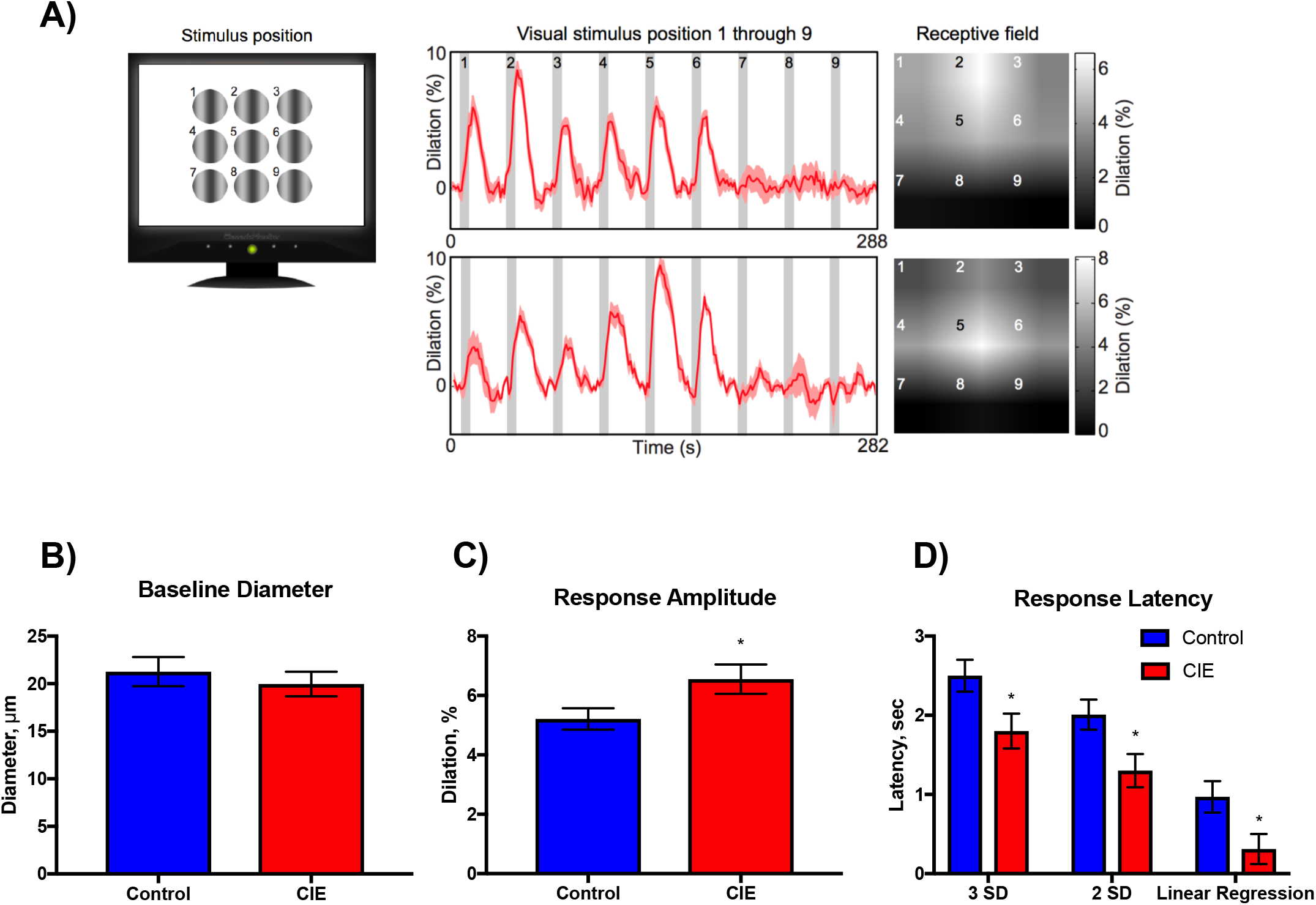
Stimulus-evoked changes in V1 blood vessel diameter. A) Cartoon of the stimulus for measuring the spatial receptive field of V1 blood vessel dilation responses and two representative examples. Time series graphs show mean (± SEM) dilation to the nine stimulus positions. Receptive field with intensity scaled to amplitude of the response at each position. Summary graphs show mean (± SEM) baseline diameter (B); stimulus-evoked dilation (C) and latency to dilate (D) for control and CIE treated mice. Symbol (*): p < 0.05, unpaired t-test.

To determine whether CIE exposure induced similar changes in vessel responses in other cortical sensory areas, a separate group of mice went through the CIE protocol and stimulus-evoked changes in vessel diameter were monitored in the barrel cortex. In Air mice, the mean diameter of vessels in the barrel cortex was similar to that seen in V1 (compare Fig. 2B vs Fig. 3A) and at baseline, there was no difference in barrel cortex vessel diameter between Air and CIE treated mice (Figure 3A). Air puffs delivered to the whiskers of head-fixed mice increased vessel diameter in Air mice by approximately 12% (Figure 3B), nearly twice that measured in V1 (mean difference 6.53 ± 1.41; unpaired t-test, t=4.62, df=133, p<0.0001). However, unlike V1 vessels, sensory-evoked dilation of barrel cortex vessels did not differ between Air and CIE groups. In Air control mice, the response latency (to 2 standard deviations of the baseline) of the dilation of barrel cortex vessels was significantly shorter than that for V1 vessels (unpaired t-test, t=14.98, p<0.0001), but there was no difference in the latency of barrel cortex vessel dilation between Air and CIE groups (Figure 3C). As baseline vessel diameter may influence the size of the evoked dilation, we compared the size of barrel cortex vessels with response amplitude and latency for CIE and Air exposed mice. As shown in Figure 3D, the correlations between baseline diameter and stimulus-evoked dilation or latency did not differ between groups.

**Figure 3.**
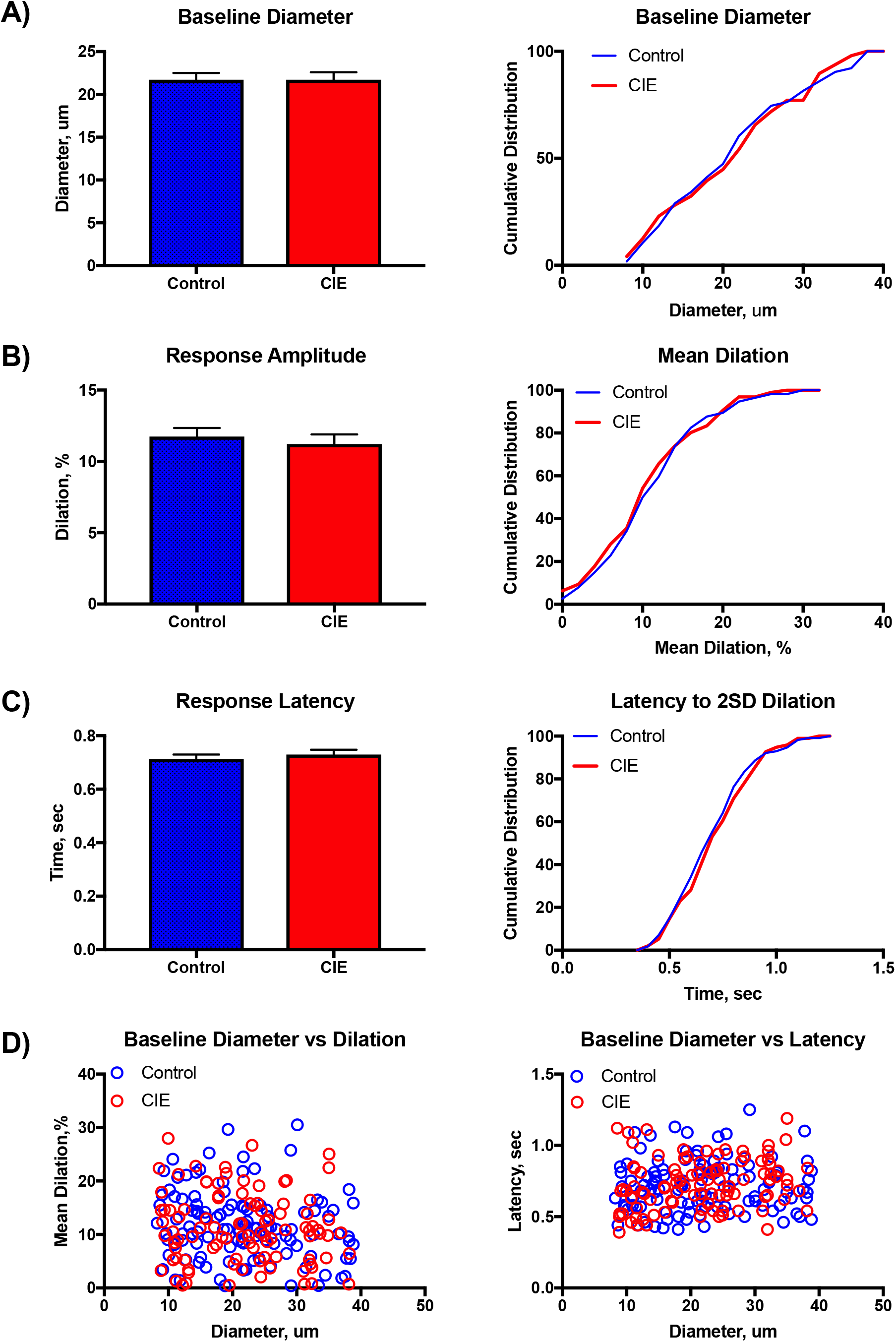
Stimulus-evoked changes in barrel cortex blood vessel diameter. Summary graphs (left column) and cumulative histograms (right column) show mean (± SEM) baseline diameter (A), stimulus-evoked dilation (B), and latency to respond (C) for control and CIE treated mice. D) Scatter plots show correlation between baseline diameter and evoked dilation (left) and response latency (right).(2013). Substance Abuse and Mental Health Administration. Results from the 2013 National Survey on Drug Use and Health: Summary of National Findings. pp NSDUH Series H-48, HHS Publication No. (SMA) 14-4863.

## Discussion

The findings of the present study show that repeated cycles of CIE exposure produce differential effects on sensory-evoked neural and vessel responses in mice. CIE treated mice showed alterations in the amplitude and orientation index of V1 neurons compared to control animals while the rate and magnitude of vessel dilation following stimulation was enhanced. The effect on vessel dynamics was selective for V1 as sensory evoked changes in vessel diameter and latency in barrel cortex were not different between Air and CIE treated mice. To our knowledge, this is the first study to use in vivo two-photon imaging to assess the effects of chronic ethanol exposure on neural and vessel responses. As this CIE treatment protocol induces dependence in mice that is accompanied by enhanced ethanol consumption (den Hartog *et al*, 2016; Lopez *et al*, 2005), these results suggest that humans with alcohol use disorders may show aberrant neural and vessel responses that are sensory modality dependent.

A wealth of studies using slice electrophysiology report that repeated exposures of rodents to alcohol induces changes in the excitability and synaptic properties of neurons in frontal cortex (Holmes *et al*, 2012; Kroener *et al*, 2012; Nimitvilai *et al*, 2016; Salling *et al*, 2018) and sub-cortical (Brodie, 2002; Hopf *et al*, 2007; Varodayan *et al*, 2017; Wang *et al*, 2007; Williams *et al*, 2018) areas. Fewer studies have examined the effects of ethanol on neuronal responses in visual or somatosensory cortices and these have largely focused on acute or neonatal exposures (reviewed by Medina and Krahe, 2008). For example, an early report by Begleiter and colleagues showed that repeated intubations of adult rats with ethanol resulted in enhanced visual-evoked potentials during withdrawal that persisted in the visual cortex for more than a month (Begleiter and Porjesz, 1977). Pertubations in visual-evoked responses were also observed in rats fed an ethanol containing liquid diet although these changes partially recovered following a week of abstinence (Kjellstrom *et al*, 1994). Neuroimaging studies in humans report mixed effects during sensory evoked responses. In one study, alcohol dependent subjects who had been recently detoxified showed reduced occipital cortex activation during presentation of a checkerboard visual cue as compared to healthy controls (Hermann *et al*, 2007). In contrast, a recent meta-analysis reported greater activation of visual cortex in alcohol-dependent subjects as compared to controls when subjects were presented with alcohol versus neutral cues (reviewed by Hanlon *et al*, 2014). Similar results were observed following delivery of drug-specific images for individuals with substance-use disorders. These findings suggest that while alcohol exposure may dampen neuronal responses to irrelevant or neutral cues, it may enhance those associated with its use consistent with the idea of attentional bias.

In the present study, responses of V1 neurons to the drifting grating stimuli was slightly reduced in CIE exposed mice and this was accompanied by a modest increase in the orientation selectivity. Paradoxically, in these same animals, the rate and amplitude of changes in V1 arteriole diameter were enhanced in the CIE treated animals. These data suggest impaired coupling between neural and vascular responses in CIE exposed mice. Using a video imaging approach, Mayhan and colleagues performed a series of studies examining the effects of a liquid ethanol diet on vessel responses in the parietal cortex of rats. Baseline diameter was different between control and ethanol-fed rats but dilation in response to a variety of compounds including isoproterenol, histamine, cromalkin (agonist of ATP-gated potassium channels) was reduced in ethanol exposed animals rats while no change was noted for nitroglycerin (Mayhan, 1992; Mayhan and Didion, 1996). This group also reported that ethanol-fed rats had altered vessel responses following inhibition or activation of inwardly rectifying potassium (KiR) channels (Sun *et al*, 2008). Related to this finding are a number of reports (reviewed by Cannady *et al*, 2018) showing changes in the expression of function of a variety of potassium channels including those activated by G-proteins (GIRK) and calcium (SK2 and BK).

While we observed region-dependent differences in the effects of CIE on vascular responses in the present study, it is important to note that studies in the visual cortex were done with anesthetized mice while those in the barrel cortex used awake mice acclimated to head fixation. This along with differences in the type and/or intensity of stimulation used to evoke responses may have contributed to the observed effects. Follow-up studies using multi-region imaging are thus necessary to validate this hypothesis.

## Acknowledgments

The authors thank Dr. Marcelo Lopez for assistance with the ethanol vapor studies. This project was funded by NIH grants R21AA022168 (JJW), R37AA009986 (JJW) and P50AA010761.

